# White matter alterations in autism spectrum disorder and attention-deficit/hyperactivity disorder in relation to sensory profile

**DOI:** 10.1101/656264

**Authors:** Haruhisa Ohta, Yuta Aoki, Takashi Itahashi, Chieko Kanai, Junya Fujino, Motoaki Nakamura, Nobumasa Kato, Ryu-ichiro Hashimoto

## Abstract

**Background:** Autism spectrum disorder (ASD) and attention deficit hyperactivity disorder (ADHD) have high rates of co-occurrence and share atypical behavioral characteristics, including sensory problems. The present diffusion tensor imaging (DTI) study was conducted to examine whether and how white matter abnormalities are observed in adult populations with developmental disabilities (DD) and to determine how brain-sensory relationships are either shared between or distinct to ASD and ADHD.

**Methods:** We collected DTI data from adult developmental disorder (DD) populations (a primary diagnosis of ASD: n=105, ADHD: n=55) as well as age and sex matched typically developed (TD) participants (n=58). Voxel-wise fractional anisotropy (FA), mean diffusivity, axial diffusivity, and radial diffusivity (RD) were analyzed using tract-based spatial statistics. The severities of sensory problems were assessed using the Adolescent/Adult Sensory Profile (AASP).

**Results:** Categorical analyses identified voxel clusters showing significant effects of DD on FA and RD in the posterior portion of the corpus callosum and its extension in the right hemisphere. Furthermore, regression analyses using the AASP scores revealed that slopes in relationships of FA or RD with the degree of sensory problems were parallel between the two DDs in large parts of the affected corpus callosum regions, although a small but significant cluster did exist showing interaction between the diagnosis of DD and an AASP subscale score on RD.

**Conclusions:** These results indicate that white matter abnormalities and their relationships to sensory problems are largely shared between ASD and ADHD, with localized abnormalities showing significant between-diagnosis differences within DD. (247 words)

## Introduction

Autism spectrum disorder (ASD) is a developmental disorder characterized by impairment of social interaction and repeated restricted behavior (American Psychiatric Association 2013). Attention-deficit/hyperactivity disorder (ADHD) is also a developmental disorder with symptoms including attention-related difficulties and hyperactivity (American Psychiatric Association 2013). Despite the differences in their core symptoms, more than 50% of people with ASD have clinical ADHD symptoms (Joshi *et al.* 2017, Llanes *et al.* 2018), while 20-30% of people with ADHD present with clinically significant symptoms of ASD (Grzadzinski *et al.* 2011, Grzadzinski *et al.* 2016). Furthermore, family members of individuals with one disorder are at risk of developing not only the one but also the other syndrome (Sandin *et al.* 2014, Chen *et al.* 2017, Miller *et al.* 2018). Such overlaps in symptoms and familial cross-aggregation have raised questions regarding similarity and distinction between these developmental disorders.

Neuroimaging studies contrasting either ASD or ADHD against typically developing people (TD) have shown that both disorders are characterized by abnormal connectivity in function as well as structure (reviewed in (Anagnostou and Taylor 2011, Aoki *et al.* 2013a, Castellanos and Aoki 2016, Aoki *et al.* 2018)). Thus, some prior studies using diffusion tensor imaging (DTI) have enrolled the three groups (ASD, ADHD, and TD) and reported on similarities and distinctions in white matter between the disorders compared with TD (Ameis *et al.* 2016, Aoki *et al.* 2017, Chiang *et al.* 2017). Results of categorical analyses contrasting the three diagnostic groups vary, possibly because of heterogeneity of the developmental disorders (Pelphrey *et al.* 2011, Ecker and Murphy 2014). However, they consistently emphasize the corpus callosum (Ameis *et al.* 2016, Aoki *et al.* 2017, Chiang *et al.* 2017). Besides categorical analysis, dimensional analyses were also conducted in these studies in which all the participants were allocated to one group and the relationship between DTI parameters and ASD symptoms was examined. Brain-ASD symptom relationships were reported in dimensional analyses across the diagnostic groups, suggesting people with ASD and with ADHD shared these relationships.

Sensory symptoms include both hyper- and hypoactivity to textures, smelling, touching, visual, or auditory input (reviewed in (Robertson and Baron-Cohen 2017)). Practically, sensory symptoms are one of the diagnostic criteria of ASD, but not for ADHD in DSM-5 (American Psychiatric Association 2013). Indeed, earlier studies reported that sensory symptoms were evident in more than 90% of people with ASD (Leekam *et al.* 2007), while not being seen in many cases of ADHD (Reynolds and Lane 2009). However, more recent studies observed atypical sensory profiles in both pediatric (Ghanizadeh 2011) and adult populations (Bijlenga *et al.* 2017), indicating that individuals with ADHD might suffer from sensory problems, perhaps to a lesser extent than ASD. Given that sensation is an input of external stimuli beginning at birth, sensory problems could underlie development of impaired social interaction (Orefice *et al.* 2016). In fact, sensory problems may cascade into higher-order dysfunction in individuals with ASD (Robertson and Baron-Cohen 2017, Thye *et al.* 2018).

As mentioned above, prior studies have examined the brain-symptom relationship across different clinical diagnoses with the perspective of ASD symptoms (Ameis *et al.* 2016, Aoki *et al.* 2017, Chiang *et al.* 2017). However, to the best of our knowledge, no study has examined the similarity or distinction of the components of social deficits between individuals with ASD and with ADHD. Given that in individuals with ASD, sensory problems may underlie the development of ASD symptoms (reviewed in (Robertson and Baron-Cohen 2017)), investigation of relationships between sensory problems and the brain across diagnostic groups would deepen our understanding of whether ASD symptoms observed in individuals with ASD and with ADHD share the same roots.

The aim of the current study was three-fold. First, we examined the effect of a diagnosis of ASD and ADHD on DTI parameters in 218 adults with or without developmental disorders to capture consistencies or inconsistencies in the results of prior studies. Then, we performed dimensional analyses to examine similarities in the brain-sensory symptoms relationship across diagnostic groups. Finally, we conducted interaction analyses to see distinctions in brain-sensory symptoms relationships between diagnostic groups. We selected data with small levels of head motion because it impacts the results of DTI analysis (Yendiki *et al.* 2014). We used the Adolescent/Adult Sensory Profile (AASP) to assess sensory symptoms (Brown and Dunn 2002).

## Method

### Participants

We recruited 160 adults with a primary diagnosis of ASD (*n* = 105) or ADHD (*n* = 55) and 58 TD participants, matched for age and sex. After a multidisciplinary team, consisting of psychiatrists and psychologists, assessed all participants, clinical diagnosis of ASD and ADHD were made based on DSM-IV-TR. Among the 105 individuals with ASD, we administered the Autism Diagnostic Observation Schedule (ADOS) (Gotham *et al.* 2007, Gotham *et al.* 2009) to 83 individuals. All participants in the ASD group who underwent the ADOS satisfied the diagnostic criteria for ASD. To exclude the comorbidity of ASD from ADHD, the ADOS was carried out in 21 out of 55 subjects in the ADHD group. Only one of the participants with ADHD met the diagnostic criteria for ASD using ADOS. The diagnosis of ADHD has been confirmed by administering Conners’ Adult ADHD Diagnostic Interview for DSM-IV (CAADID) in all of the 55 participants in ADHD group (Epstein, Johnson, & Conners, 2001). To further characterize participants, the Autistic Spectrum Quotient (AQ) was obtained from 192 participants (ASD: *n* = 101, ADHD: *n* = 33, TD: *n* = 58) (Baron-Cohen *et al.* 2001). Data for assessment of ADHD severity was obtained from 150 participants (ASD: *n* = 86, ADHD: *n* = 46, TD: *n* = 18) using Conner’s Adult ADHD Rating Scales (CAARS) (Conners *et al.* 1999). The intelligence quotient (IQ) scores of the clinical participants (ASD: *n* = 105, ADHD: *n* = 44) were evaluated using either the Wechsler Adult Intelligence Scale-Third Edition (WAIS-III) or WAIS-Revised (WAIS-R) (Wechsler 1981, Wechsler 1997). TD participants were recruited through advertisements or acquaintances. Lack of a psychiatric diagnosis in TD participants was confirmed using the Structured Clinical Interview for DSM-IV Axis I Disorders (DATA 1997). In the TD group, IQ scores were estimated using a Japanese version of the National Adult Reading Test (JART) (Matsuoka *et al.* 2006). Twelve participants were taking antipsychotics (10 ASD, 2 ADHD), while thirty-four participants had been administered stimulants (32 ADHD, 2 ASD). Exclusion criteria for all the participants included any history of head trauma, serious medical or surgical illness, or substance abuse. All the participants were confirmed to have a full-scale IQ above 74.

Sensory symptoms were evaluated using the subscale of AASP (Brown and Dunn 2002) (ASD: *n* = 62, ADHD: *n* = 44, TD: *n* = 38). The AASP is a self-reported questionnaire consisting of 60 items from the following sensory sections; taste/smell processing, movement processing, visual processing, touch processing, activity level, and auditory processing. Participants were asked to respond to each item on a five-point Likert scale from “almost never” to “almost always”. Each item belongs to one of four subscales: Low Registration (hyposensitivity), Sensation Seeking, Sensory Sensitivity (hypersensitivity), and Sensation Avoiding. One participant failed to complete all the items included for Sensation Avoiding. These subscales were contrasted in the three groups using *F*-tests. To correct multiple comparisons, we adopted the Bonferroni method and set the threshold for significance at *P* < 0.0125 (=0.05/4: the number of subscales). The institutional review board of Showa University Karasuyama Hospital approved all of the procedures adopted in this study. Written informed consent was obtained from all the participants after fully explaining the purpose of this study. The authors assert that all procedures contributing to this work comply with the ethical standards of the relevant national and institutional committees on human experimentation and with the Helsinki Declaration of 1975, as revised in 2008.

### Data acquisition

All magnetic resonance imaging (MRI) data were obtained using a 3T MR Scanner (MAGNETON Verio; Siemens Medical Systems, Erlangen, Germany) with a 12-channel head coil. Diffusion-weighted images were acquired using a single-shot, spin-echo, echo planar imaging sequence. The acquisition parameters were as follows: repetition time = 10,300 ms, echo time = 102 ms, field of view=200 × 200 mm, 75 contiguous axial slices of 2.0-mm thickness without gap, phase-encoding direction = anterior-posterior, 65 non-collinear motion-probing gradients, and *b* = 1000 s/mm^2^. The directions of gradients were optimized according to a previous study (Caruyer *et al.* 2013). The acquisition of the images included ten images without diffusion-weighting (*b0*) interspersed throughout the sequence.

### Preprocessing

Images were preprocessed with FSL version 5.0 (FMRIB Software Library, http://www.fmrib.ox.ac.uk). DTI data are potentially at risk for a wide variety of artifacts, including motion artifacts and eddy current. Therefore, automatic artifact correction was conducted for all images using DTIPrep (Oguz *et al.* 2014). Susceptibility induced distortion was corrected in all acquired images using TOPUP implemented in FSL (Andersson *et al.* 2003, Smith *et al.* 2004). After the DTIPrep and TOPUP pipelines were performed, all data were registered to the first *b*=0 image with affine transformation for correcting distortions. We used FSL rmsdiff functions to calculate root-mean-square (RMS) deviation of absolute intervolume displacement with respect to the first image of each run (Jenkinson *et al.* 2012). Because the results of comparisons of DTI parameters are particularly sensitive to head motion (Yendiki *et al.* 2014), participants with a mean RMS over 5 mm were excluded from this study (Aoki *et al.* 2017).

### Tract-Based Spatial Statistics (TBSS) preprocessing

The images were then skull-stripped and factional anisotropy (FA), mean diffusivity (MD), axial diffusivity (AD), and radial diffusivity (RD) images were calculated using the DTIFIT function for all subjects. Generated FA images were registered to the Montreal Neurological Institute (MNI) 152 standard space using non-linear registration. Normalized FA images were averaged to create a mean FA image, which was then thinned to create a mean FA skeleton (Smith *et al.* 2006). Voxel-wise analyses using general linear models were conducted in skeleton areas with an FA of at least 0.2. Other DTI parameters, including MD, RD, and AD, were projected onto the mean FA skeleton.

### DTI group analyses

We performed an F-test to examine the main effect of diagnosis on DTI parameters using the FSL randomise tool with age, sex, and motion as nuisance covariates. The contrasts were tested with 5,000 permutations. The statistical threshold was defined at *P* < 0.05, correcting for multiple comparisons by threshold-free cluster enhancement (TFCE). We focused on clusters with a minimum size of 10 voxels. Post-hoc pairwise group comparisons were conducted for clusters with a significant main effect of diagnosis on DTI parameters.

### DTI dimensional analysis

We performed the dimensional analyses when we examined the relationships between sensory symptoms and DTI parameters. In these analyses, we used a vector of AASP subscale scores of all the subjects as an effect of interest. Here, we performed the analysis only for FA and RD maps because a significant main effect of diagnosis was not identified in the MD or AD map (see Results). The analyses were conducted on voxels of the binary mask image identified by the *F*-tests of FA and RD. The nuisance covariates included age, sex, and motion. The analysis was performed independently for all the four AASP subscales. The statistical threshold for significance was defined at *P* < 0.05, the TFCE corrected, and the spatial extension threshold was set to *k* > 10 voxels.

### DTI interaction analysis

We examined the interaction of the slope in relationships between the sensory symptoms (Low Registration, Sensation Seeking, Sensory Sensitivity, and Sensation Avoiding) and DTI parameters of FA and RD. The variables of interest were the element-wise product of the vector of an AASP subscale and the vector representing diagnostic status: (i) ASD or non-ASD, (ii) ADHD or non-ADHD, and (iii) TD or non-TD. The three vectors representing the diagnostic status (i-iii) were also included in the model. The other nuisance covariates included age, sex, and motion. The analyses were conducted on voxels of the binary mask image identified in the F-test and repeated for all the four AASP subscales. We adopted a threshold for statistical significance at *P* < 0.05, corrected for multiple comparisons using TFCE with a minimal number of voxels larger than 10.

## Results

### Demographic data

Table 1 shows the demographic and clinical data of participants in the ASD, ADHD, and TD groups. *F*-tests showed no significant differences in age, sex, or full scale IQ (FIQ) among the three groups (*P* > 0.1). The main effect of diagnosis was observed in all of the four subscales of the AASP. Post-hoc tests of Low Registration, Sensory Sensitivity, and Sensation Avoiding showed the same pattern when compared with TD; individuals with ASD and those with ADHD showed significantly higher scores, with no significant difference between these two clinical groups (Table 1). On the other hand, Sensation Seeking exhibited a different pattern in which individuals with ASD showed significantly lower scores compared to both the TD group and individuals with ADHD, which turned out not to be significantly different from each other.

### DTI categorical analyses

The categorical analyses with FA showed the main effect of diagnosis on two clusters in the corpus callosum (Table 2 and Figure 1-A). Post-hoc analyses revealed that individuals with ASD and with ADHD have significantly smaller FA values compared with TD, whereas ASD and ADHD were comparable (Figure 1-B). The categorical analysis with RD also showed a main effect of diagnosis in the right posterior part of the corpus callosum (Table 2 and Figure 1-C). Post-hoc tests using RD values extracted from the two clusters showed that, compared with TD participants, individuals with ASD and with ADHD had statistically significantly larger RD values, whereas there was no significant difference between ASD and ADHD (Figure 1-D). However, we found a localized significant cluster where ASD showed larger RD than ADHD. Other DTI parameters, such as MD and AD, did not show any significant main effect of diagnosis.

**Figure 1.**
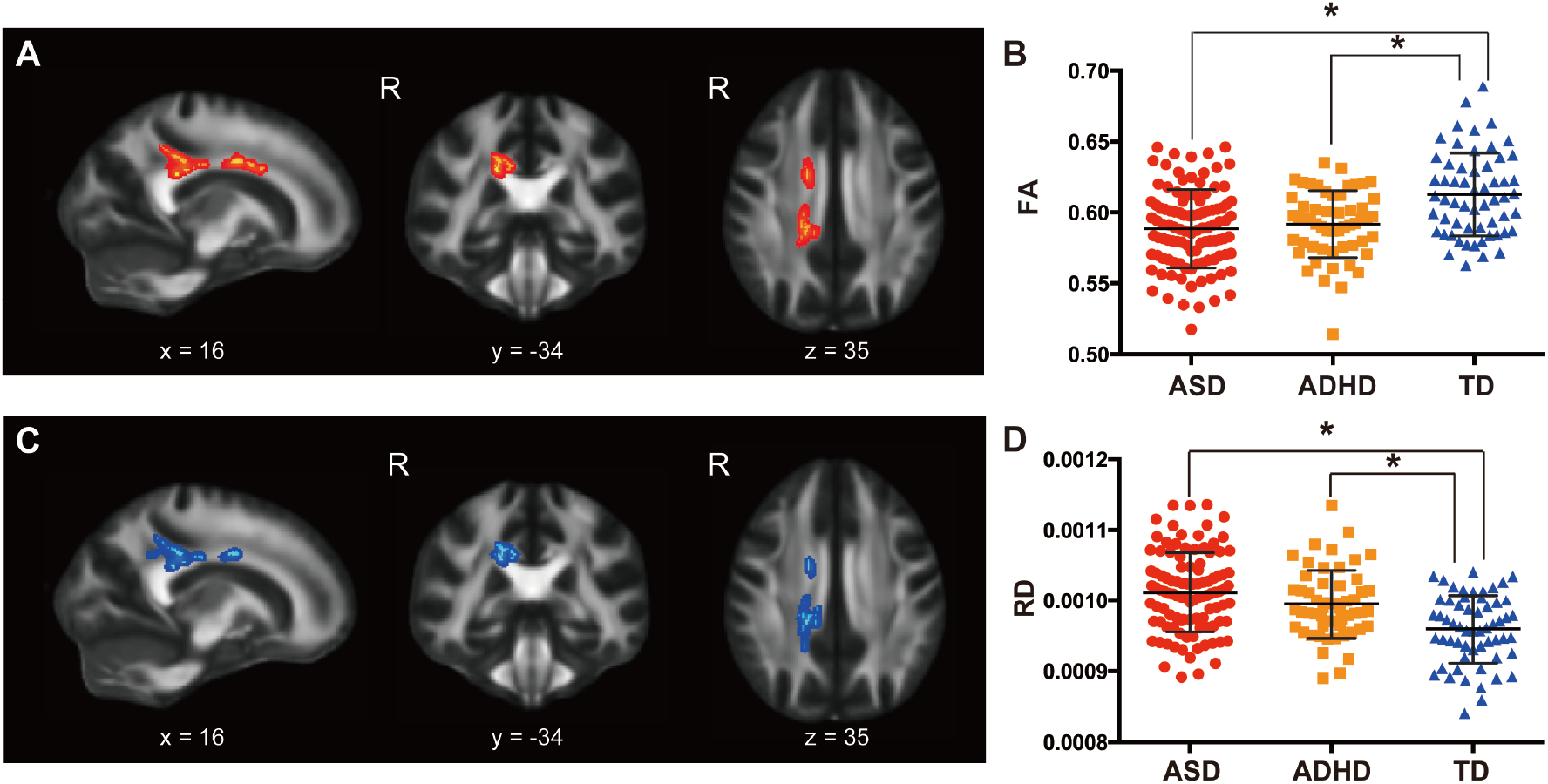
Significant main effect of diagnosis of developmental disorders on fractional anisotropy (FA) and radial diffusivity (RD). (A) Significant clusters of voxels showing a main effect of diagnosis on FA. (B) Plots of mean FA values extracted from significant voxels shown in (A). (C) Significant clusters of voxels showing a main effect of diagnosis on RD. Note the high extent of spatial overlapping with (A). (D) Plots of mean RD values extracted from significant voxels shown in (C). The asterisk (*) indicates a significant difference between groups (*P* < 0.05).

### DTI dimensional analyses

The dimensional analyses revealed that RD values in the posterior body of the corpus callosum were negatively associated with Sensation Seeking (Table 3 and Figure 2-A, -B) and positively associated with Sensation Avoiding (Table 3 and Figure 2-C, -D). We extracted mean RD values from identified voxels and confirmed that slopes in relationships of the AASP subscale with the DTI parameter were comparable among the three groups (*F*(2, 138) = 1.185, *P* = 0.31 for Figure 2B; *F*(2, 137) = 0.036, *P* = 0.96 for Figure 2D). The analysis with FA showed that the DTI parameter in the posterior body of the corpus callosum was negatively correlated with Sensory Sensitivity (Figure 2-E, F) and Sensation Avoiding (Table 3 and Figure 2-G and H). Slopes of the AASP subscale – FA relationships were comparable among the three groups (*F*(2, 138) = 0.90, *P* = 0.41 for Figure 2F; *F*(2, 138) = 0.22, *P* = 0.80 for Figure 2G).

**Figure 2.**
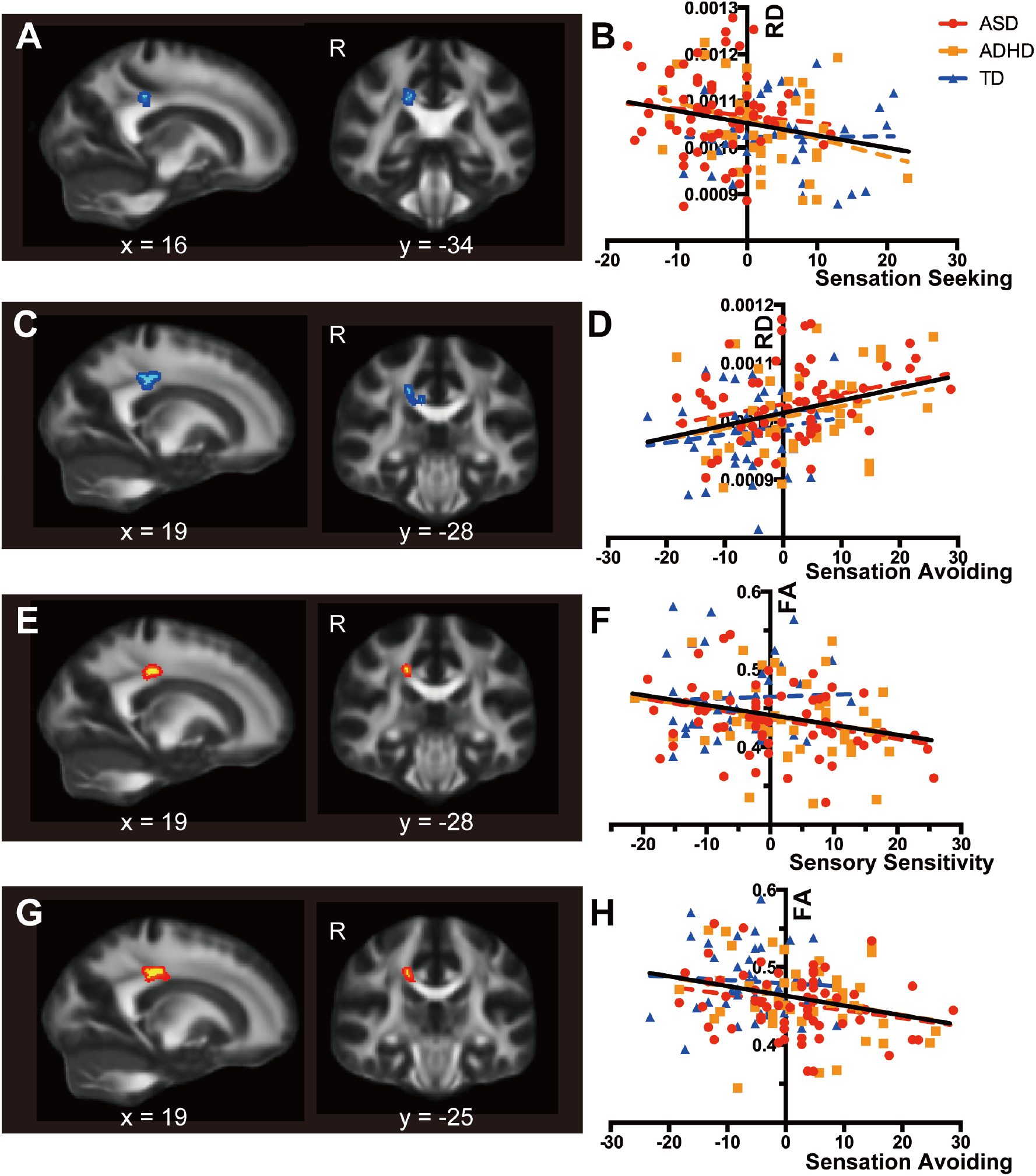
Significant voxels identified by dimensional analyses using subscale scores of the sensory profile. (A) Significant voxels identified by regression of radial diffusivity (RD) on the Sensation Seeking score. For the sake of visualization, the voxel clusters were thickened using the tbss_fill script implemented in FSL. (B) Scatterplots and regression lines showing relationships between the demeaned Sensation Seeking score and RD values extracted from voxels shown in (A). Colored dotted lines indicate regression lines for the data of autism spectrum disorder (red), attention-deficit/hyperactivity disorder (orange), and typically developed participants (blue), whereas the black lines indicate regression lines for the combined data of the three groups. (C) Significant voxels identified by regression of RD on the Sensation Avoiding score. (D) Scatterplots and regression lines showing relationships between the demeaned Sensation Avoiding score and RD values. RD values were extracted from significant voxels in (C). (E) Significant voxels identified by regression of FA on the Sensory Sensitivity score. (F) Scatterplots and regression lines showing relationships between the demeaned Sensation Sensitivity score and FA values extracted from voxels shown in (E). (G) Significant voxels identified by regression of FA on the Sensation Avoiding score. (H) Scatterplots and regression lines showing relationships between the demeaned Sensation Avoiding score and FA values extracted from voxels shown in (G).

### DTI interaction analyses

Significant interaction in the Sensory Sensitivity–FA slope was observed between TD and DD groups. People with ASD had negative correlation between Sensory Sensitivity score and FA in the midbody of the corpus callosum, while TD people showed positive correlation (Table 3 and Figure 3-A, -B). An *F*-test confirmed significant group differences in the slopes of the FA-Sensory Sensitivity relationship (*F*(2, 138) = 5.57, *P* = 0.005). On the other hand, the analysis with RD showed significant interaction in Sensory Sensitivity score and RD value between ASD and ADHD groups (Table 3 and Figure 3-C, -D). Individuals with ASD showed positive correlation between RD value and Sensory Sensitivity scores, while individuals with ADHD had a negative correlation. We observed significant group differences in the slopes of the RD-Sensation Sensitivity relationship (*F*(2, 138) = 9.61, *P* = 0.0001). The cluster was located in the right posterior corpus callosum. Other subscales of the AASP did not show any significant results in any DTI parameter.

**Figure 3.**
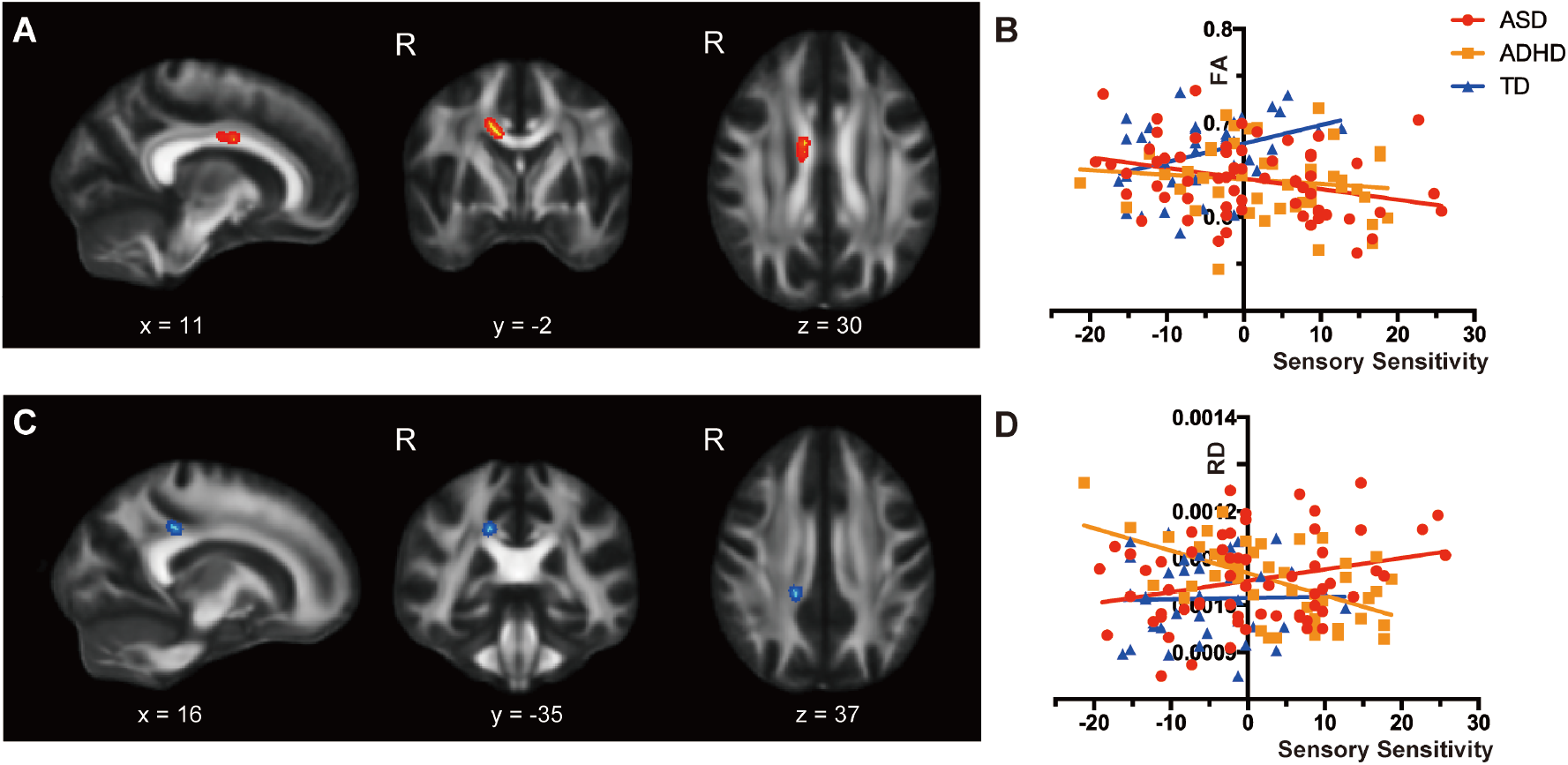
Significant voxels identified by interaction analyses of diagnosis status and subscale scores of sensory profile. (A) Significant voxels identified by the contrast between developmental disorders (autism spectrum disorder [ASD] and attention-deficit/hyperactivity disorder [ADHD]) and typically developed participants for the regression of fractional anisotropy (FA) on the Sensory Sensitivity score. For the sake of visualization, the voxel clusters were thickened using the tbss_fill script implemented in FSL. (B) Scatterplots and regression lines showing relationships between the demeaned Sensory Sensitivity score and FA values extracted from voxels in (A). (C) Significant voxels identified by the contrast between ASD and ADHD for the regression of radial diffusivity (RD) on the Sensory Avoiding score. (D) Scatterplots and regression lines showing relationships between the demeaned Sensation Avoiding score and RD values extracted from voxels in (C).

## Discussion

The current TBSS study enrolled a relatively large number of participants: 218 adults (TD: *n* = 58, ASD: *n* = 105, ADHD: *n* = 55). In the conventional approach of categorical group comparisons, we used *F*-tests and identified the main effects of diagnosis on FA and RD values in regions around the body and the splenium of the corpus callosum. Post-hoc tests revealed that, compared with the TD group, FA values were smaller in individuals with ASD and with ADHD, while they were not significantly different from each other. In terms of RD, individuals with ASD and with ADHD had larger values compared with the TD group, again, in the posterior part of the corpus callosum. The dimensional analysis, using the scores of sensory profiles, demonstrated the brain regions where the three groups showed comparable relationships between the DTI parameters and sensory problems. In contrast, the interaction analyses showed significant results in other brain regions; the DD groups had a negative association between FA and Sensory Sensitivity, while the TD group showed a positive correlation. The interaction analysis with RD showed that individuals with ASD and those with ADHD had different associations between white matter organization and Sensory Sensitivity. These findings demonstrated both similarities and distinctions in DTI parameters and in relationships with sensory problems between the two developmental disorders of ASD and ADHD.

Previously, two studies directly contrasted three groups using TBSS (Ameis *et al.* 2016, Aoki *et al.* 2017). Strikingly, Ameis et al. identified smaller FA values in the posterior part of the corpus callosum in individuals with ASD and with ADHD compared with the TD group, and their results were anatomically close to our current results (Ameis *et al.* 2016). In contrast, while the results showed astonishing anatomical overlap with the current study, another study reported different patterns (Aoki *et al.* 2017). This study showed smaller FA values in individuals with ASD compared with individuals with ADHD and TD. Given that DTI values differ between children, adolescents, and adults (Qiu *et al.* 2008), one potential explanation for inconsistency in the ADHD results is the difference in age. Aoki et al. recruited children aged between 6 and 13 (Aoki *et al.* 2017). Our current study enrolled adults aged between 20 and 55, while the age of the participants in the Ameis et al. study is between the age range of these two studies. Future research that fully depicts the developmental trajectory of DTI parameters in each brain region is expected to better examine this question.

The present study participants with ADHD exhibited severe sensory problems to a degree comparable to, or even higher than, individuals with ASD in the three subscales (Low Registration, Sensory Sensitivity, and Sensation Avoiding). Although the results were not consistent with our expectations, they were fully consistent with the results from a study contrasting the scores of Sensory Profile (Dunn 1999) across people with ASD, ADHD, or TD, except for the scores for Sensation Seeking (Little *et al.* 2018). While the current study showed a significant difference in Sensation Seeking between individuals with ASD and with ADHD, Little et al. found no significant difference between them. Clinical differences in the participants could be one possible reason for the inconsistency. Little et al. did not perform a semi-structured diagnostic interview nor provide any information on medication status. Given that ASD and ADHD often co-occur, it would be important to make a maximum effort to differentiate ASD from ADHD for the comparison of the two groups. Medication status may also impact results, and this status was not recorded in the study authored by Little et al. For example, neurotransmitters, such as GABA, are involved in sensory processing (Cohen Kadosh *et al.* 2015, Robertson and Baron-Cohen 2017). Given that neurotransmitter levels are altered in individuals with developmental disorders (Aoki *et al.* 2012, Aoki *et al.* 2013b) and that medication influences neurotransmitter levels, medication may also influence sensory problems. As there is no established pharmacological intervention for sensory problems in developmental disorders (Case-Smith *et al.* 2015), future studies are expected to explore novel approaches to treat sensory problems in developmental disorders.

The dimensional analysis identified the body of the corpus callosum as being associated with sensory problems. Given that the region involves fibers connecting sensory areas of both hemispheres (Hofer and Frahm 2006), it is reasonable to say that the anatomical location of the current results may correlate with sensory problems. It was striking that the best fit line in the analyses for individuals with ASD, with ADHD, and TD participants were quasi-parallel. These findings suggest that these brain regions were related to sensory symptoms, regardless of the clinical diagnosis. Given the possibility that sensory symptoms contribute to the development of ASD symptoms (Robertson and Baron-Cohen 2017), the similarity in the brain-sensation symptoms relationship suggests that the process of developing ASD traits is shared by all three groups. However, it should be noted that similarity was claimed on the basis of there being no significant differences in the slopes of regression of a DTI parameter on a sensory profile score, not on the basis of rejecting a null hypothesis that the three lines differed from each other.

The analysis showed significant interaction between Sensory Sensitivity and FA values in the midbody of the corpus callosum between individuals with ASD and the TD controls. Although it did not reach statistical significance, individuals with ADHD showed a different association between Sensory Sensitivity and FA values compared with TD, and the relationship was similar to that of individuals with ASD. These results suggest that the brain region identified by the current analysis is associated with pathological brain-symptoms relationships, regardless of the clinical diagnosis (i.e. ASD or ADHD). Although the cluster was small, the anatomical location was close to the regions in which prior studies had identified the main effect of diagnosis (Ameis *et al.* 2016, Aoki *et al.* 2017). We speculate that the difference in sensory problems in participants might have contributed to inconsistency in the results pertaining to this brain area across these studies (Ameis *et al.* 2016, Aoki *et al.* 2017).

There are some limitations in the current study. First, the age range of the participants is wide (from 20 to 55 y). As we included age as a covariate of nuisance and there was no significant difference in age between groups, the impact of this wide age range was minimized. In addition, although wide age range may increase the heterogeneity of participants, few studies enrolled people with ASD in their forties or fifties, resulting in a lack of data for middle-aged individuals with ASD. The current study may contribute to filling in this gap. Second, we included both sexes. As we assumed that sex impacts the results (Baron-Cohen *et al.* 2005), we included sex as a covariate of nuisance. Reflecting male predominant prevalence, the majority of our participants was male. However, the sample size was not large enough to repeat the analysis including only male or female subjects. Future studies with large sample sizes that include only one sex are needed to test the replicability of our findings.

## Conclusion

The current DTI study enrolled adults whose primary diagnosis was ASD or ADHD and compared them to TD people to investigate differences in white matter organization around the middle to posterior parts of the corpus callosum across diagnostic groups. We investigated brain-behavior relationships from the perspective of sensory symptoms. This study showed that, in some brain regions, the three groups showed similar relationships of DTI parameters to sensory problems, while in other brain regions the groups showed different relationships between the DTI and sensory problems. The current study provided insight into similarities and distinctions in the process of development of clinical ASD symptoms and subclinical traits across ASD, ADHD, and TD.

## Conflict of interest

None

We express our gratitude to Mr. Taku Sato for his support in recruiting participants and data collection. This study was supported by the Brain Mapping by Integrated Neurotechnologies for Disease Studies (Brain/MINDS) from Japan Agency for Medical Research and Development, AMED to RH, the KAKENHI from the Japan Society for the Promotion of Science (18K15493) to YA, and the KAKENHI from the Japan Society for the Promotion of Science (15K09843) to HO.

## References

Ameis, S. H., Lerch, J. P., Taylor, M. J., Lee, W., Viviano, J. D., Pipitone, J., Nazeri, A., Croarkin, P. E., Voineskos, A. N., Lai, M. C., Crosbie, J., Brian, J., Soreni, N., Schachar, R., Szatmari, P., Arnold, P. D. and Anagnostou, E. (2016) ’A diffusion tensor imaging study in children with ADHD, autism spectrum disorder, OCD, and matched controls: distinct and non-distinct white matter disruption and dimensional brain-behavior relationships’, Am J Psychiatry, 173(12), 1213–1222.

American Psychiatric Association (2013) Diagnostic and statistical manual of mental disorders.

Anagnostou, E. and Taylor, M. J. (2011) ’Review of neuroimaging in autism spectrum disorders: what have we learned and where we go from here’, Mol Autism, 2(1), 4.

Andersson, J. L., Skare, S. and Ashburner, J. (2003) ’How to correct susceptibility distortions in spin-echo echo-planar images: application to diffusion tensor imaging.’, Neuroimage, 20(2), 870–88.

Aoki, Y., Abe, O., Nippashi, Y. and Yamasue, H. (2013a) ’Comparison of white matter integrity between autism spectrum disorder subjects and typically developing individuals: a meta-analysis of diffusion tensor imaging tractography studies’, Mol Autism, 4(1), 25.

Aoki, Y., Cortese, S. and Castellanos, F. X. (2018) ’Diffusion tensor imaging studies of attention-deficit/hyperactivity disorder: meta-analyses and reflections on head motion’, J Child Psychol Psychiatry, 59(3), 193–202.

Aoki, Y., Inokuchi, R., Suwa, H. and Aoki, A. (2013b) ’Age-related change of neurochemical abnormality in attention-deficit hyperactivity disorder: a meta-analysis’, Neurosci Biobehav Rev, 37(8), 1692–701.

Aoki, Y., Kasai, K. and Yamasue, H. (2012) ’Age-related change in brain metabolite abnormalities in autism: a meta-analysis of proton magnetic resonance spectroscopy studies’, Transl Psychiatry, 2, e69.

Aoki, Y., Yoncheva, Y. N., Chen, B., Nath, T., Sharp, D., Lazar, M., Velasco, P., Milham, M. P. and Di Martino, A. (2017) ’Association of white matter structure with autism spectrum disorder and attention-deficit/hyperactivity disorder’, JAMA Psychiatry, 74(11), 1120–1128.

Baron-Cohen, S., Knickmeyer, R. C. and Belmonte, M. K. (2005) ’Sex differences in the brain: implications for explaining autism’, Science, 310(5749), 819–23.

Baron-Cohen, S., Wheelwright, S., Skinner, R., Martin, J. and Clubley, E. (2001) ’The autism-spectrum quotient (AQ): evidence from Asperger syndrome/high-functioning autism, males and females, scientists and mathematicians.’, J Autism Dev Disord, 31(1), 5–17.

Bijlenga, D., Tjon-Ka-Jie, J. Y. M., Schuijers, F. and Kooij, J. J. S. (2017) ’Atypical sensory profiles as core features of adult ADHD, irrespective of autistic symptoms’, Eur Psychiatry, 43, 51–57.

Brown, C. and Dunn, W. (2002) Adolescent-adult sensory profile: user’s manual., San Antonio: Therapy Skill Builders.

Caruyer, E., Lenglet, C., Sapiro, G. and Deriche, R. (2013) ’Design of multishell sampling schemes with uniform coverage in diffusion MRI’, Magn Reson Med, 69(6), 1534–40.

Case-Smith, J., Weaver, L. L. and Fristad, M. A. (2015) ’A systematic review of sensory processing interventions for children with autism spectrum disorders’, Autism, 19(2), 133–48.

Castellanos, F. X. and Aoki, Y. (2016) ’Intrinsic functional connectivity in attention-deficit/hyperactivity disorder: a science in development’, Biol Psychiatry Cogn Neurosci Neuroimaging, 1(3), 253–261.

Chen, Q., Brikell, I., Lichtenstein, P., Serlachius, E., Kuja-Halkola, R., Sandin, S. and Larsson, H. (2017) ’Familial aggregation of attention-deficit/hyperactivity disorder’, J Child Psychol Psychiatry, 58(3), 231–239.

Chiang, H. L., Chen, Y. J., Lin, H. Y., Tseng, W. I. and Gau, S. S. (2017) ’Disorder-specific alteration in white matter structural property in adults with autism spectrum disorder relative to adults with ADHD and adult controls’, Hum Brain Mapp, 38(1), 384-395.

Cohen Kadosh, K., Krause, B., King, A. J., Near, J. and Cohen Kadosh, R. (2015) ’Linking GABA and glutamate levels to cognitive skill acquisition during development’, Hum Brain Mapp, 36(11), 4334–45.

Conners, C. K., Erhardt, D. and Sparrow, E. (1999) CAARS: Conner’s Adult ADHD Rating Scales., Multi-Health Systems Incorporated.

Data, D. (1997) Structured clinical interview for DSM-IV axis I disorders., Washington: American Psychiatric Press.

Dunn, W. (1999) Sensory profile: User’s manual, San Antonio, TX: Psychological Corporation.

Ecker, C. and Murphy, D. (2014) ’Neuroimaging in autism--from basic science to translational research’, Nat Rev Neurol, 10(2), 82–91.

Ghanizadeh, A. (2011) ’Sensory processing problems in children with ADHD, a systematic review’, Psychiatry Investig, 8(2), 89–94.

Gotham, K., Pickles, A. and Lord, C. (2009) ’Standardizing ADOS scores for a measure of severity in autism spectrum disorders’, J Autism Dev Disord, 39(5), 693–705.

Gotham, K., Risi, S., Pickles, A. and Lord, C. (2007) ’The Autism Diagnostic Observation Schedule: revised algorithms for improved diagnostic validity’, J Autism Dev Disord, 37(4), 613–27.

Grzadzinski, R., Di Martino, A., Brady, E., Mairena, M. A., O’Neale, M., Petkova, E., Lord, C. and Castellanos, F. X. (2011) ’Examining autistic traits in children with ADHD: does the autism spectrum extend to ADHD?’, J Autism Dev Disord, 41(9), 1178–91.

Grzadzinski, R., Dick, C., Lord, C. and Bishop, S. (2016) ’Parent-reported and clinician-observed autism spectrum disorder (ASD) symptoms in children with attention deficit/hyperactivity disorder (ADHD): implications for practice under DSM-5’, Mol Autism, 7, 7.

Hofer, S. and Frahm, J. (2006) ’Topography of the human corpus callosum revisited-comprehensive fiber tractography using diffusion tensor magnetic resonance imaging’, Neuroimage, 32(3), 989-94.

Jenkinson, M., Beckmann, C. F., Behrens, T. E., Woolrich, M. W. and Smith, S. M. (2012) ’Fsl’, Neuroimage, 62(2), 782–90.

Joshi, G., Faraone, S. V., Wozniak, J., Tarko, L., Fried, R., Galdo, M., Furtak, S. L. and Biederman, J. (2017) ’Symptom profile of ADHD in youth with high-functioning autism spectrum disorder: a comparative study in psychiatrically referred populations’, J Atten Disord, 21(10), 846–855.

Leekam, S. R., Nieto, C., Libby, S. J., Wing, L. and Gould, J. (2007) ’Describing the sensory abnormalities of children and adults with autism’, J Autism Dev Disord, 37(5), 894–910.

Little, L. M., Dean, E., Tomchek, S. and Dunn, W. (2018) ’Sensory processing patterns in autism, attention deficit hyperactivity disorder, and typical development’, Phys Occup Ther Pediatr, 38(3), 243–254.

Llanes, E., Blacher, J., Stavropoulos, K. and Eisenhower, A. (2018) ’Parent and teacher reports of comorbid anxiety and ADHD symptoms in children with ASD’, J Autism Dev Disord, 2018/08/01.

Matsuoka, K., Uno, M., Kasai, K., Koyama, K. and Kim, Y. (2006) ’Estimation of premorbid IQ in individuals with Alzheimer’s disease using Japanese ideographic script (Kanji) compound words: Japanese version of National Adult Reading Test’, Psychiatry Clin Neurosci, 60(3), 332–9.

Miller, M., Musser, E. D., Young, G. S., Olson, B., Steiner, R. D. and Nigg, J. T. (2018) ’Sibling recurrence risk and cross-aggregation of attention-deficit/hyperactivity disorder and autism spectrum disorder’, JAMA Pediatr, 2018/12/12.

Oguz, I., Farzinfar, M., Matsui, J., Budin, F., Liu, Z., Gerig, G., Johnson, H. J. and Styner, M. (2014) ’DTIPrep: quality control of diffusion-weighted images’, Front Neuroinform, 8, 4.

Orefice, L. L., Zimmerman, A. L., Chirila, A. M., Sleboda, S. J., Head, J. P. and Ginty, D. D. (2016) ’Peripheral mechanosensory neuron dysfunction underlies tactile and behavioral deficits in mouse models of ASDs’, Cell, 166(2), 299-313.

Pelphrey, K. A., Shultz, S., Hudac, C. M. and Vander Wyk, B. C. (2011) ’Constraining heterogeneity: the social brain and its development in autism spectrum disorder’, J Child Psychol Psychiatry, 52(6), 631–44.

Qiu, D., Tan, L. H., Zhou, K. and Khong, P. L. (2008) ’Diffusion tensor imaging of normal white matter maturation from late childhood to young adulthood: voxel-wise evaluation of mean diffusivity, fractional anisotropy, radial and axial diffusivities, and correlation with reading development’, Neuroimage, 41(2), 223-32.

Reynolds, S. and Lane, S. J. (2009) ’Sensory overresponsivity and anxiety in children with ADHD.’, Am J Occup Ther, 63(4), 433–40.

Robertson, C. E. and Baron-Cohen, S. (2017) ’Sensory perception in autism’, Nat Rev Neurosci, 18(11), 671–684.

Sandin, S., Lichtenstein, P., Kuja-Halkola, R., Larsson, H., Hultman, C. M. and Reichenberg, A. (2014) ’The familial risk of autism’, JAMA, 311(17).

Smith, S. M., Jenkinson, M., Johansen-Berg, H., Rueckert, D., Nichols, T. E., Mackay, C. E., Watkins, K. E., Ciccarelli, O., Cader, M. Z., Matthews, P. M. and Behrens, T. E. (2006) ’Tract-based spatial statistics: voxelwise analysis of multi-subject diffusion data’, Neuroimage, 31(4), 1487–505.

Smith, S. M., Jenkinson, M., Woolrich, M. W., Beckmann, C. F., Behrens, T. E., Johansen-Berg, H., Bannister, P. R., De Luca, M., Drobnjak, I., Flitney, D. E., Niazy, R. K., Saunders, J., Vickers, J., Zhang, Y., De Stefano, N., Brady, J. M. and Matthews, P. M. (2004) ’Advances in functional and structural MR image analysis and implementation as FSL’, Neuroimage, 23 Suppl 1, S208-19.

Thye, M. D., Bednarz, H. M., Herringshaw, A. J., Sartin, E. B. and Kana, R. K. (2018) ’The impact of atypical sensory processing on social impairments in autism spectrum disorder’, Dev Cogn Neurosci, 29, 151–167.

Wechsler, D. (1981) WAIS-R manual: Wechsler adult intelligence scale-revised.

Wechsler, D. (1997) Wechsler adult intelligence scale-III., San Antonio, TX: Pearson

Yendiki, A., Koldewyn, K., Kakunoori, S., Kanwisher, N. and Fischl, B. (2014) ’Spurious group differences due to head motion in a diffusion MRI study’, Neuroimage, 88, 79–90.

